# Reversible hypervascularization drives cognitive decline and blood-brain barrier damage during aging

**DOI:** 10.64898/2026.05.04.720441

**Authors:** Mihail Ivilinov Todorov, Katalin Todorov-Völgyi, David-Paul Minde, Saketh Kapoor, Mayar Ali, Rainer Malik, Johannes Christian Paetzold, Julian McGinnis, Lanyue Zhang, Alessia Nottebrock, Lu Liu, Mikael Simons, Harsharan Singh Bhatia, Marios K. Georgakis, Martin Dichgans, Farida Hellal, Ali Ertürk

## Abstract

Cerebrovascular dysfunction emerges early in neurodegeneration, yet how vascular structure, blood–brain barrier (BBB) failure, and cognition are linked remains undefined. Using VesselPro, a whole-brain pipeline integrating perfusion-resolved 3D imaging with spatial proteomics, we mapped vascular aging across the mouse lifespan. We identify two discrete trajectories: a hypovascular state and a previously unrecognized hyper-vascular, BBB-compromised state, defined relative to a young-adult vascular baseline. The hyper-vascular trajectory, concentrated in the cortex and hippocampus, was associated with marked spatial memory impairment and pervasive BBB leakage. Spatial proteomics revealed a coordinated program involving angiogenic activation, endothelial stress, cytoskeletal remodeling, and inflammatory signaling. Cross-species comparison with human proteomic biomarkers from the UK Biobank showed strong alignment between the mouse hypervascular signature and vascular dementia risk, but minimal concordance with Alzheimer’s disease, defining a vascular-specific dementia endotype. Transient Tie2 activation with AKB-9778 attenuated this trajectory, improving vessel organization, BBB integrity, and memory performance. Our findings show that endothelial instability is a key mechanistic driver of this heterogeneous vascular aging state.

**Graphical abstract:** 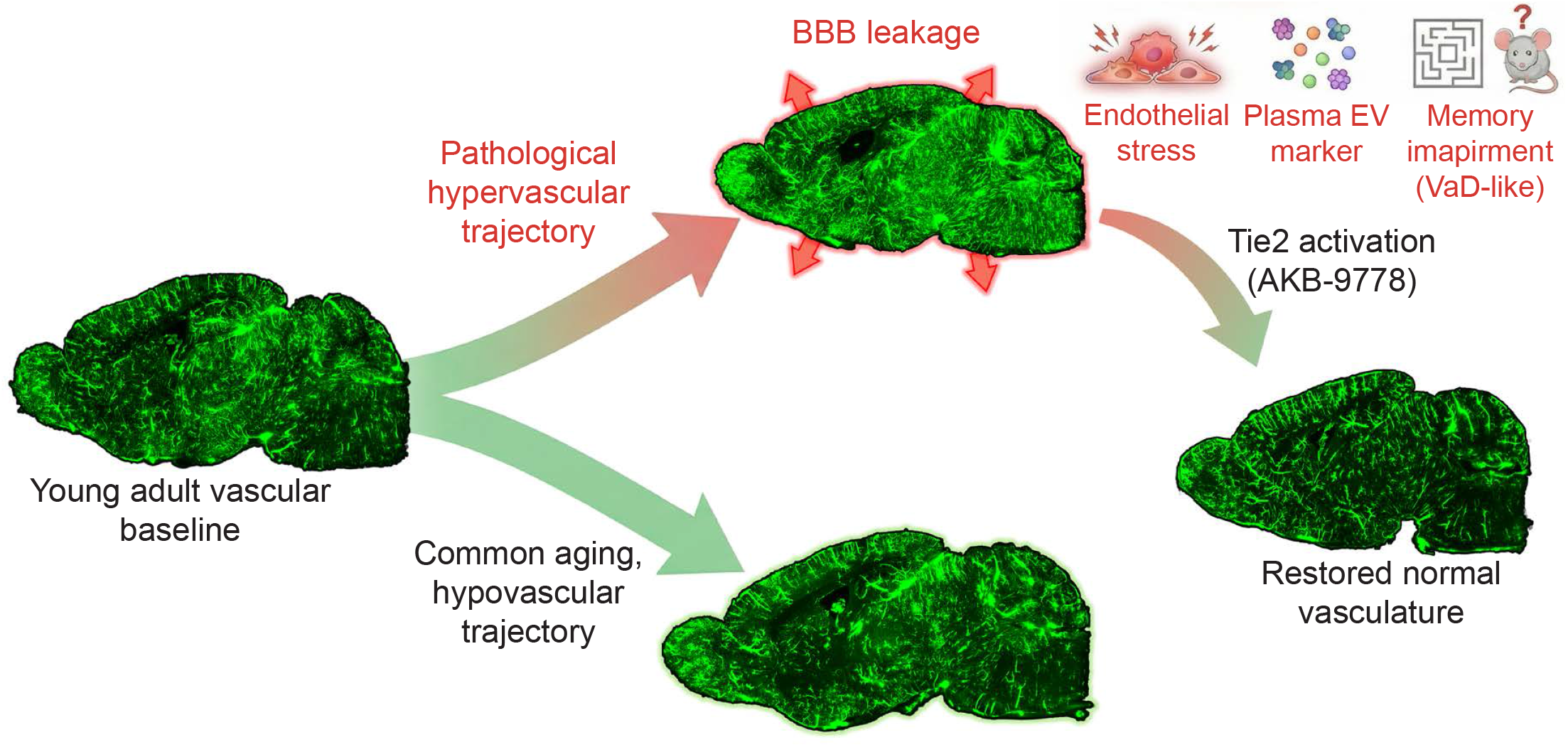

**Highlights:**

- **VesselPro**: we developed a pipeline for **capturing perfused blood vessels** in the brain: unexpected bifurcation in vascular aging: traditional hypovascular state and a novel, **hypervascular, BBB-compromised state**.
- **Hypervascular pathology**: Linked the hypervascular profile in the cortex and hippocampus to **spatial memory impairment, pervasive BBB leakage**, and **angiogenic inflammatory signaling**.
- **Clinical endotype**: Cross-species analysis via the **UK Biobank** confirms this hypervascular signature aligns with **vascular dementia** risk rather than Alzheimer’s disease.
- **Pharmacological stabilization**: Demonstrated that **Tie2 signaling activation (AKB-9778)** improves the vessel organization, BBB integrity, and memory performance.

## Introduction

Cerebrovascular dysfunction is a primary driver of vascular cognitive impairment and vascular dementia (VaD), contributing directly to cognitive decline independent of amyloid-β plaques and tau neurofibrillary tangles^1,2^. One of the earliest detectable vascular changes is disruption of the blood-brain barrier (BBB), the endothelial interface that regulates molecular exchange between the blood and neural tissue^3,4^. Human imaging, biomarker, and post-mortem studies demonstrate BBB leakage in cerebral small vessel disease, mild vascular cognitive impairment, and VaD, including in cognitively normal individuals with high vascular risk^5,6^. BBB breakdown correlates with endothelial dysfunction, pericyte loss, and markers of small vessel disease^7,8^ and predicts cognitive decline^9^, establishing BBB disruption as a central pathological mechanism and emerging biomarker in vascular dementia. Paradoxically, early cerebrovascular dysfunction can present with regional hyperperfusion. MRI and PET studies have reported increased flow in hippocampal and temporal circuits in AD and MCI^10–12^, as well as across vascular-driven brain pathologies, such as vascular cognitive impairment and cerebral small vessel disease, where capillary dysfunction, flow heterogeneity, and blood–brain barrier impairment are central features. These findings, also replicated in experimental models of aging and vascular perturbation^13^, suggest that increased perfusion and vascular remodeling represent a recurring feature of human vascular brain pathology, although their structural basis, molecular regulation, and functional consequences remain poorly understood. Such hyperperfusion may initially reflect compensation^14^ but can enhance vascular stress and promote inflammation and BBB leakage, accelerating neurodegeneration^15^. Age-related vascular pathology also includes aberrant remodeling^16–18^. While angiogenesis may restore perfusion^19^, newly formed vessels often remain structurally immature and leaky^20^. Increased vessel density is a typical response to hypoxic states (28–30) and is observed in vulnerable hippocampal regions in AD^21,22^. Vascular endothelial growth factor (VEGF) driven angiogenic signaling is frequently elevated in hypoxia and neuroinflammatory disease^23–26^ and can compromise BBB integrity by weakening tight junctions, increasing paracellular permeability, and altering endothelial-cytoskeletal coupling^27–34^. These observations suggest that cerebrovascular aging may involve heterogeneous remodeling processes that are not fully captured by average trends alone. Notably, in young adulthood, cerebrovascular organization is characterized by a tightly regulated, homeostatic vascular architecture with low inter-individual variability. Aging may therefore not simply shift this baseline uniformly, but destabilize it, allowing divergent vascular outcomes to emerge across individuals. However, existing methods cannot resolve these potential trajectories. Macroscopic neuroimaging modalities lack either sufficient resolution or adequate brain coverage^35^, while microCT casting^36^ prevents analysis of barrier permeability and molecular states. Optical clearing enables 3D vascular imaging^37–40^ but typically requires blood removal and cannot reliably retain BBB tracers; blood-retaining variants^41^ have not been applied to aging, where extracellular matrix dysregulation^42^ further complicates staining. As a result, no current approach can simultaneously map perfused microvasculature, BBB integrity, and molecular remodeling across the entire brain. To overcome these limitations, we developed VesselPro, a whole-brain pipeline combining perfusion-resolved (vessels with active circulation) 3D vascular imaging, quantitative BBB mapping, and spatial proteomics within the same specimens. By integrating perfusion-resolved vascular measurements with structural metrics and molecular profiling under standardized clearing and imaging conditions, this approach minimizes technical variability. It enables discrimination between biological vascular states and processing-related artifacts, providing a scalable and reproducible framework for systematic analysis of cerebrovascular aging. This integrated framework enables unbiased quantification of inter-individual vascular variability, allowing us to determine whether aging-associated heterogeneity reflects a uniform continuum around a preserved physiological baseline, or instead arises from breakdown of a shared young-adult vascular state into discrete, mechanistically distinct phenotypes with different cognitive consequences, and to test causal links between vascular state, BBB integrity, and cognitive function through targeted endothelial stabilization.

## Results

### A brain-wide pipeline for quantifying perfused vasculature, blood-brain barrier integrity, and spatial proteomic remodeling in the aging mouse brain

To capture brain-wide structural and functional changes in the aging vasculature, we established a multimodal pipeline optimized for aging brain analysis. This workflow integrates unbiased mapping of functionally perfused blood vessels^43^, optical tissue clearing and spatial proteomics^44^, enabling comprehensive structural and molecular profiling from the same cohort of adult and aged C57BL/6 mice (6-30 months) within a single experimental framework. VesselPro (**Fig. 1a**) combines high-resolution 3D light sheet imaging, automated vessel quantification, BBB leakage mapping (**Fig. 1a’**), and region-specific proteomics to directly link vascular architecture within in situ molecular signatures. Mice were injected intraperitoneally with Evans Blue (EB) to assess BBB permeability and subsequently underwent intracardiac injection of wheat germ agglutinin conjugated to Alexa Fluor 594 to label actively perfused vessels. Brains were harvested, optically cleared, and imaged using fluorescent light sheet microscopy to generate volumetric maps of the entire functional brain vasculature. This approach provided high-resolution visualization of the complete network of perfused vessels, including capillaries, across the whole brain, yielding a fixed single-time-point snapshot of vascular perfusion for post-mortem analysis (**Fig. 1b**). Selective quantification of perfused vessels at whole brain scale was achieved, which is not possible with standard post-fixation staining. Automated analysis of the 3D datasets yielded vessel length, bifurcation density, and topological features for each brain and region. Following imaging, the same brains were rehydrated and anatomically dissected according to 3D imaging results. Spatial proteomic profiles were generated via the DISCO-MS workflow for regions identified as having abnormal perfused vascular length or BBB leakage. This integrated design enabled direct comparison of perfused vascular networks (**Fig. 1c**), BBB integrity, and corresponding regional proteomes within the same mice, providing a comprehensive view of structural, functional, and molecular aging signatures across the brain.

**Fig.1:**
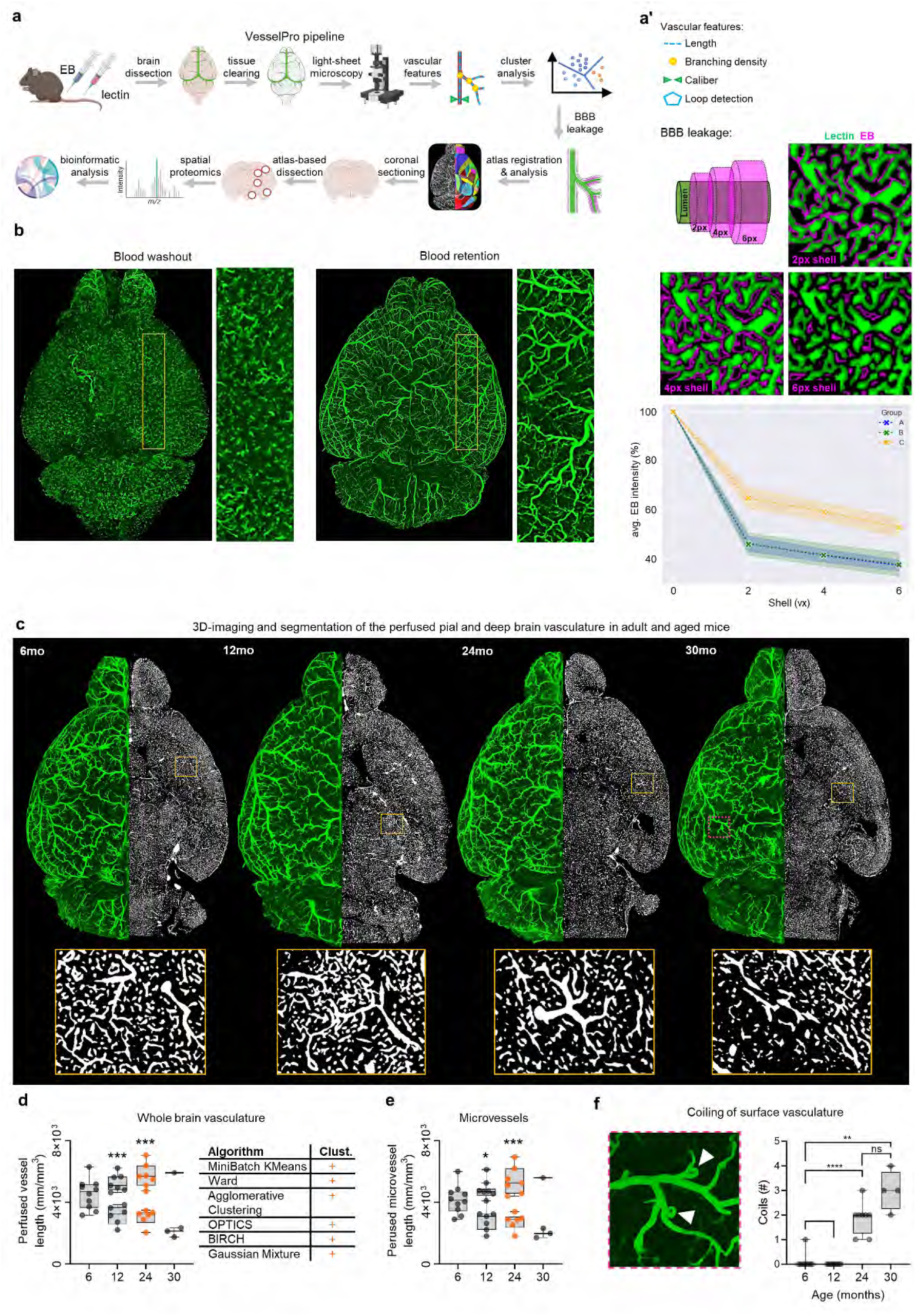
3D light-sheet microscopy imaging of whole brain functional vasculature. **a:** Schematic overview of VesselPro, a streamlined pipeline integrating capture of perfused vasculature in adult mice, optical tissue clearing, high-resolution imaging of perfused vessels and extravasation tracers, automated vessel segmentation with quantitative blood– brain barrier (BBB) leakage mapping, and spatial proteomics for region-specific molecular profiling, enabling direct correlation of vascular architecture with in situ proteomic signatures. **a’:** Insert of (a) parameters quantified to systematically characterize the perfused brain vascular network (vessel length, bifurcation density, caliber, microvascular loop count, distance-resolved average Evans blue intensity). **b:** 3D visualization of the cerebrovascular network in 12 months old mice after blood washout or blood retention which enables a snapshot of brain perfusion state. **c:** 3D visualizations (green) and single plane of the segmented perfusion maps (white) at 6, 12, 24 and 30 months of age. **d:** Quantification of perfused vascular length in the brains at 24-month of age reveals two distinct aging trajectories: classical vascular rarefaction or hypervascularization. Unsupervised clustering (Agglomerative Clustering, BIRCH, Gaussian Mixture Models, MiniBatch KMeans, OPTICS, and Ward’s method) robustly separated these two phenotypes, mostly contributed by (**e**) the microvessels. **f:** High-resolution imaging of age-related vascular alterations on the brain surface: coiling of large surface vessels near cortical entry points. The box and whisker plot shows the median, data spread (whiskers), and individual data points.

### Aged mice of identical age diverge into hypovascular and hypervascular brain phenotypes

Using the brain-wide perfusion maps generated with VesselPro, we identified pronounced regional and age-dependent increases in inter-individual variability in perfused vascular length in the 12- and 24-month-old cohorts. The 6-month-old mice exhibited a narrow, unimodal distribution of vascular length across brain regions, indicating a stable and homogeneous young-adult vascular baseline. Rather than forming a continuous distribution around the young-adult reference, aged animals clustered toward two distinct extremes characterized by either reduced or elevated perfused vascular length (**Fig. 1d**) driven by the microvasculature (defined as vessels with one to three voxel radii, ≤15 µm) (**Fig. 1e**). The subgroup with reduced vessel length was consistent with classical vascular rarefaction. In contrast, the second subgroup exhibited markedly increased vascular length. To objectively assess this heterogeneity, we applied multiple unsupervised clustering methods (Agglomerative Clustering, BIRCH, Gaussian Mixture Models, Mini-Batch KMeans, OPTICS, Ward’s method), all of which consistently separated the 24-months-old aged brains into two reproducible groups based on brain-wide perfused vascular length (**Fig. 1d**), without evidence for a stable intermediate vascular state in aged mice These results demonstrate that aging is associated with the loss of the young-adult vascular homeostasis, resulting in substantial heterogeneity in cerebrovascular organization among mice of identical chronological age. In addition, high-resolution imaging revealed coiling of large superficial vessels near cortical branch points before penetrating the parenchyma in aged animals (**Fig. 1f**); the functional significance of this morphological feature remains unknown.

### Hypervascular brains exhibit regionally localized vascular remodeling

To identify the anatomical basis of the vascular heterogeneity observed among 24-month-old animals, we performed a region-specific analysis comparing both vessel length and bifurcation density to those of 6-month-old (6mo) brains, which exhibited a uniform vascularization profile at the whole-brain level. In the 24-month-old subgroup with reduced vascularization (24mo-Lo) compared to the 6-months-old group, we identified 19 brain regions with a consistent and statistically significant (p < 0.05) decrease in perfused vessel length, predominantly localized to subcortical structures, hypothalamic nuclei, and the cerebellum (**Fig. 2a**). Parallel analysis of the vascular bifurcation density patterns revealed that 15 of these 19 regions also displayed significantly reduced bifurcation density (**Fig. 2b**), indicating a coordinated loss of both perfused vessel abundance and network complexity in these areas. Conversely, the highly vascularized 24-month-old subgroup (24mo-Hi) exhibited widespread, robust increases in both vessel length and bifurcation density across 34 distinct brain regions, primarily within the cerebral cortex, hippocampal formation, and cerebellum (**Fig. 2c-d**). Notably, 36 regions showed elevated branching complexity, consistent with active vascular remodeling.

**Fig.2:**
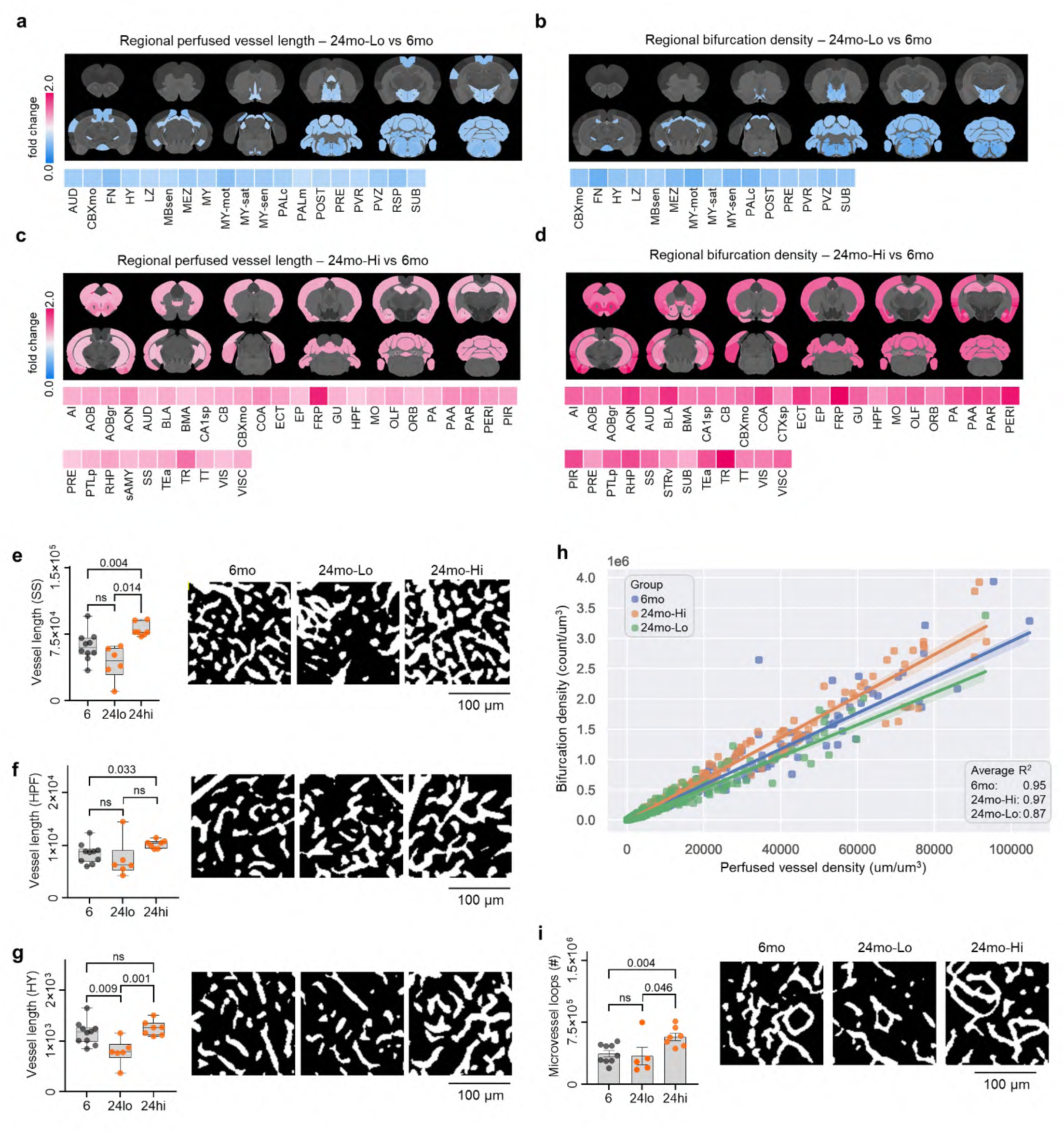
Region-specific changes in perfused vessel length, bifurcation complexity, and microvascular organization in aged mouse brains. **a**,**b:** region specific changes in perfused vessel length and bifurcation density in the hypovascularized 24 months old (24mo-Lo) versus the 6 months old (6mo). **c**,**d:** region specific changes in perfused vessel length and bifurcation density in the hypervascularized 24 months old (24mo-Hi) versus the 6 months old (6mo). **e-g:** Quantitative comparisons and vascular network representations in the somatosensory cortex (SS), hippocampal formation (HPF), and hypothalamus (HY) illustrate subgroup-specific differences in topology. **h:** Robust linear regression analysis of length and bifurcation density in the perfused vasculature of 6mo and 24-month-old subgroups. Each point represents the R^2^ regression coefficient for the group’s median in a given brain region. **i:** Quantification of closed microvascular loops revealed a significant increase in the 24mo-Hi compared to both the 6mo and the 24mo-Lo, suggesting altered microvascular organization associated with vascular remodeling. The bar plot shows the median, ±SEM (whiskers), and individual data points.

Representative vascularization patterns and quantitative analyses from the somatosensory cortex, hippocampal formation, and hypothalamus are shown in **Fig. 2e-g**. Robust linear regression analyses were conducted for each brain region to examine the relationship between perfused vascular length and bifurcation density across three groups: 6mo 24mo-Hi, and 24mo-Lo (**Fig. 2h**). We observed a strong coupling between these vascular parameters in the young adult group (R^2^ = 0.95), consistent with a homeostatic, well-regulated vascular architecture. Interestingly, the highest coupling was found in the 24mo-Hi (R^2^ = 0.97), indicating an even tighter relationship between vessel abundance and network complexity that may reflect increased or sustained remodeling activity within structurally constrained vascular networks. In contrast, the 24mo-Lo showed a lower correlation (R^2^ = 0.87), suggesting a relative decoupling of perfused vessel length and bifurcation density. Because uniform tissue scaling would be expected to preserve proportional relationships between vascular metrics, this altered coupling argues against global processing effects and instead points toward subgroup-specific reorganization of microvascular topology. In addition to the changes in vessel length and bifurcation density, the 24mo-Hi exhibited a higher number of closed microvascular loops than both the 6mo brains and the 24mo-Lo (**Fig. 2i**).

### Distinct molecular programs define divergent vascular phenotypes

To investigate the molecular profile associated with these vascular features, we rehydrated the brain tissue samples and we performed proteomic profiling of the whole brain and the major brain regions which showed the most significant changes in the vascular network (vessel length and bifurcation density) of 24mo-Lo or 24mo-Hi mice (**Fig. 3a**). We first asked whether vascular phenotypes are associated with global differences in brain protein composition. Principal component analysis (PCA) of the brain proteomes revealed distinct clustering of the 6mo, 24mo-Hi, and 24mo-Lo samples, with minimal overlap between the clusters, indicating substantial differences in global protein expression profiles among these groups (**Fig. 3b**). Compared to brain proteomes of 6-month-old mice, 24mo-Hi exhibited a substantially greater number of significantly dysregulated proteins than the 24mo-Lo (**Fig. 3c,d**, and **Fig. S1**). The hippocampus (HC) and somatosensory cortex (SS) were selected for in-depth molecular analysis because we detected the highest number of significantly altered proteins in these regions. Analysis of the ten most significantly altered vascular-related proteins in the HC and SS regions of 24mo-Hi and 24mo-Lo subgroups (compared to 6mo) demonstrated clear region-specific expression patterns, with no overlap between the aged subgroups, indicating that vascular topology is associated with distinct proteomic signatures across brain regions. To interpret these differences at the level of biological function, we next performed Gene Ontology (GO) enrichment analysis which revealed a coordinated network of biological processes linking vascular remodeling, neuronal development, and cellular stress responses: processes related to blood vessel development, angiogenesis, and regulation of vessel size and circulation, as well as neurogenesis, neuron projection morphogenesis, and synapse organization. GO terms also encompassed cell adhesion, migration, cytoskeletal organization, and protein phosphorylation. Enrichment of apoptotic and autophagic pathways was also observed. Quantitative comparison of enriched biological processes in both 24mo-Hi and 24mo-Lo mice revealed that high vascularization was associated with broader upregulation of 30 biological processes enriched above a ratio of 1. In contrast, the 24mo-Lo showed only five processes exceeding this threshold (**Fig. 3e**). Finally, to directly link molecular changes to vascular architecture, we examined correlations between individual protein expression levels and vascular metrics across brain regions. For each protein, correlation coefficients were calculated between abundance levels and five vascular parameters: vessel length, length of microvessels, density of bifurcations, density of bifurcations in microvessels, and average vessel radius. Strong associations were defined as |r| > 0.9. Proteins were then categorized based on the directionality of association into exclusively positive, exclusively negative, or mixed positive and negative, and depicted in circular graphs representing the distribution of protein-vascular correlations by anatomical region for 6-month-old mice and 24-month-old mice with either high or low vascularization (**Fig 3f**). Of these five parameters in young animals, the average vessel radius held the strongest relationship with protein levels in the hippocampus and somatosensory cortex. In contrast, in aged mouse brains, the correlations with the other morphometric features became more evenly distributed, with a comparable number of associations across parameters. Notably, the hippocampus of 24mo-Hi mice displayed the highest number of protein-vascular associations, nearly eightfold higher than in the hippocampus of young mice. In contrast, the SS of the 24mo-Lo exhibited the most significant number of associations. Our results reveal that aging and vascularization levels strongly influence the relationship between protein expression and vascular metrics across brain regions, with the hippocampus showing the most substantial changes in 24mo-Hi mice.

**Fig.3:**
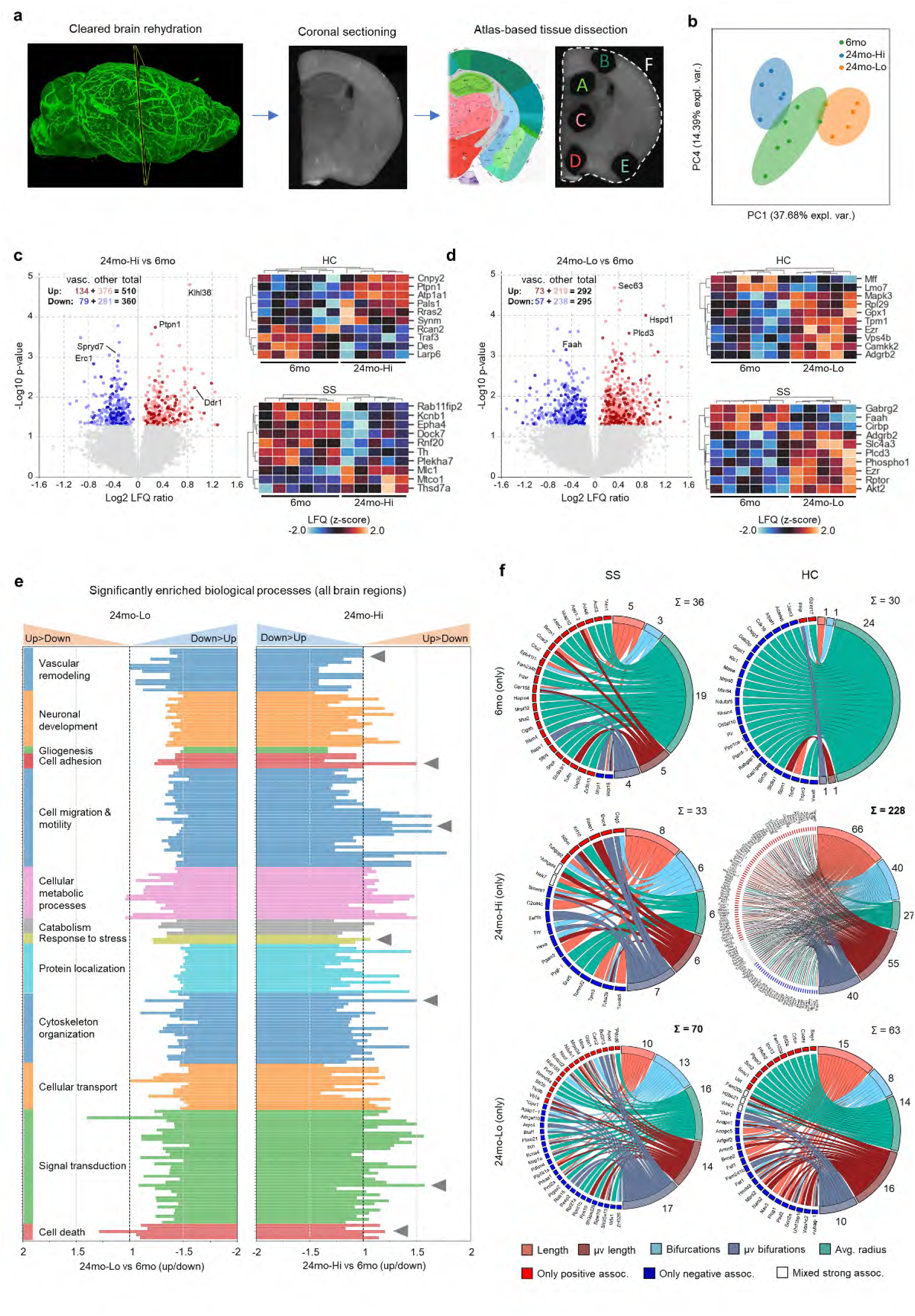
Proteomic profiling reveals distinct molecular signatures associated with cerebrovascular features in aging mouse brains. **a:** Schematic overview of the workflow used for proteomic analysis of tissue cleared brains and a representative image of the anatomical clusters extracted (A-F). **b:** Principal component analysis (PCA) of brain proteomes shows distinct clustering of 6mo, 24mo-Hi, and 24mo-Lo mice. **c, d:** Volcano plot, quantification of the differentially expressed proteins. Vascular-related proteins are shown with dark red and blue. Ten most differentially abundant proteins are ranked in the heatmap for each individual animal. **e:** Gene Ontology enrichment analysis of the differentially expressed proteins highlights molecular pathways related to vascular remodeling, neurodevelopment, and cellular stress responses, the 24mo-Hi brains exhibit substantially more pathways with upregulated protein expression. **f:** Correlations between protein expression levels (left) and vascular metrics (right) across brain regions. Circular graphs depict the normalized distribution of protein–vascular correlations by anatomical regions (HC, SS). Adjacent to each protein color indicates the level of correlations (|r| > 0.9): bright red for positive association, dark blue for negative, and white for mixed positive and negative associations.

### The hypervascular trajectory is associated with BBB leakage and spatial memory impairment

To investigate whether the observed age-related differences in vascular architecture are associated with cognitive performance, we conducted behavioral testing in 6- and 24-month-old mice. Spatial memory was assessed using the Barnes maze (BM), anxiety-related behavior was evaluated with the elevated plus maze, and motor coordination was assessed with the rotarod assay. Prior to sacrifice, blood plasma was collected for molecular profiling of extracellular vesicles (EV) and brains were processed using VesselPro (**Fig. 4a**). EV preparation circumvents the high dynamic range of bulk plasma, enriching tissue-specific signaling cargo and improving the detection sensitivity for low-abundance markers of neurovascular health^45^. While 24-month-old mice did not exhibit differences in anxiety-like behavior and showed a significant decrease in motor performance compared to 6-month-old controls (**Fig. 4b,c**), a marked divergence in spatial memory performance emerged within the aged cohort. Specifically, one subgroup of aged mice used predominantly goal-directed (targeted) search strategies in the Barnes maze, approximately twice as often as random strategies, and rapidly located the escape hole. In contrast, the other subgroup displayed impaired memory performance, characterized by reliance on random search strategies nearly threefold more frequently, and could not find the escape hole within the given time. Based on this behavioral difference, aged mice were stratified into cognitively impaired and unimpaired groups (**Fig. 4d**). Comparison of time and distance to the goal across groups revealed that the cognitively impaired aged subgroup required significantly more time and traveled a greater distance than both the 6-month-old group and the cognitively unimpaired aged subgroup (**Fig. 4e**). Importantly, quantitative analysis of the perfused brain microvessels revealed that the cognitively impaired subgroup displayed a hypervascular phenotype (**Fig. 4f**) similar to the 24mo-Hi subgroup (**Fig. 1d**). Microvessels contributed to length differences across 24 brain regions between the impaired and the unimpaired groups (**Fig. 4g**), notably within cortical and hippocampal areas, regions essential for spatial memory encoding (**Fig. 4g’**). Thus, impaired memory was associated with an increase in microvascular length across memory-relevant networks. To determine whether the hypervascular state was accompanied by BBB dysfunction, we quantified Evans Blue (EB) extravasation. Vascular masks were applied to the EB channel, and EB fluorescence intensity was quantified in concentric radial shells (2, 4, and 6 pixels) extending from perfused vessel boundaries (**Fig. 1a’**). Normalizing parenchymal to intravascular signal enabled 3D mapping of barrier integrity. These analyses showed substantial BBB leakage in the cognitively impaired subgroup. Compared to 6mo mice, cognitively impaired 24mo mice showed leakage in 50% of the analyzed brain regions. In contrast, unimpaired aged mice showed leakage in only 14% (**Fig. 4h**). The unimpaired aged subgroup had markedly elevated EB accumulation in 13 regions including the cortex, hippocampus, and thalamus, and to a lesser extent in the brainstem (**Fig. 4i**). We observed a partial overlap (six regions) between brain regions showing increased perfused microvessels and those exhibiting BBB leakage (**Fig. 4j**), demonstrating a spatial correspondence between hypervascular remodeling and barrier failure, including key regions implicated in memory processing such as the cortical and hippocampal areas (**Fig. 4i’**). To obtain orthogonal insights into the molecular composition of plasma proteomes in aging mice, we developed a simplified and scaled down version of Mag-Net^45^ to enrich negatively charged particles from plasma (comprising EVs and other membrane-bound particles), Nano-Mag-Net. Plasma EV proteomics identified 93 upregulated and 47 downregulated proteins. STRING DB indicated that 62 upregulated and 27 downregulated proteins are expressed in brain tissue (**Fig. 4k**). GO enrichment analysis revealed dysregulation in several biological processes, including cell adhesion, and innate immune response (**Fig. 4l**). Over-Representation Analysis (ORA) linked these signatures to human disease concepts (**Fig. 4m**), such as “Memory Loss” and “Forgetful”. STRING network analysis identified a densely interconnected protein cluster, with increased App, Mapk3, Tuba4a, Itm2b, Csf1r, Egfr, and decreased Agt, Mapk1, And Rac1 (**Fig. 4m’, m”**). These systemic signatures are consistent with endothelial stress and inflammatory activation accompanying the hypervascular, BBB-leaky phenotype. To evaluate the translational relevance of the plasma EV signatures associated with the hypervascular and cognitively impaired phenotype, we performed a cross-species analysis comparing our mouse data to human plasma proteomics from the UK Biobank (UKB). We examined whether proteins dysregulated in hypervascular, cognitively impaired aged mice matched human dementia-associated profiles. Using UKB Olink data, we applied Cox proportional hazard models adjusted by age and sex to identify proteins associated with All-Cause Dementia (ACD), Vascular Dementia (VaD), and Alzheimer’s Disease (AD). Analyses were restricted to proteins with sufficient case numbers. We defined a translational “match” as a protein showing both a statistically significant association in humans (p < 0.05) and concordant directionality (i.e., elevated or reduced in both species) (**Fig. 4n, pie chart**). In the ACD analysis, 65 significantly associated proteins were shared between species; of which 23 (35%) showed concordant directionality (**Fig. 4n, left**), indicating that the hypervascular mouse signature reflects broad dementia-associated patterns. The most substantial alignment was observed for VaD: of 38 significantly shared proteins, 27 (71%) exhibited matching directionality (**Fig. 4n, right**). In contrast, the overlap with AD was more limited, with only 7 of 27 (26%) significant proteins showing concordant directionality (**Fig. S2a**). Together, these findings show that the hypervascular, BBB-leaky phenotype in aged mice closely mirrors the systemic proteomic signature of human vascular dementia but not Alzheimer’s disease, identifying this trajectory as a VaD-like molecular endotype.

**Fig.4:**
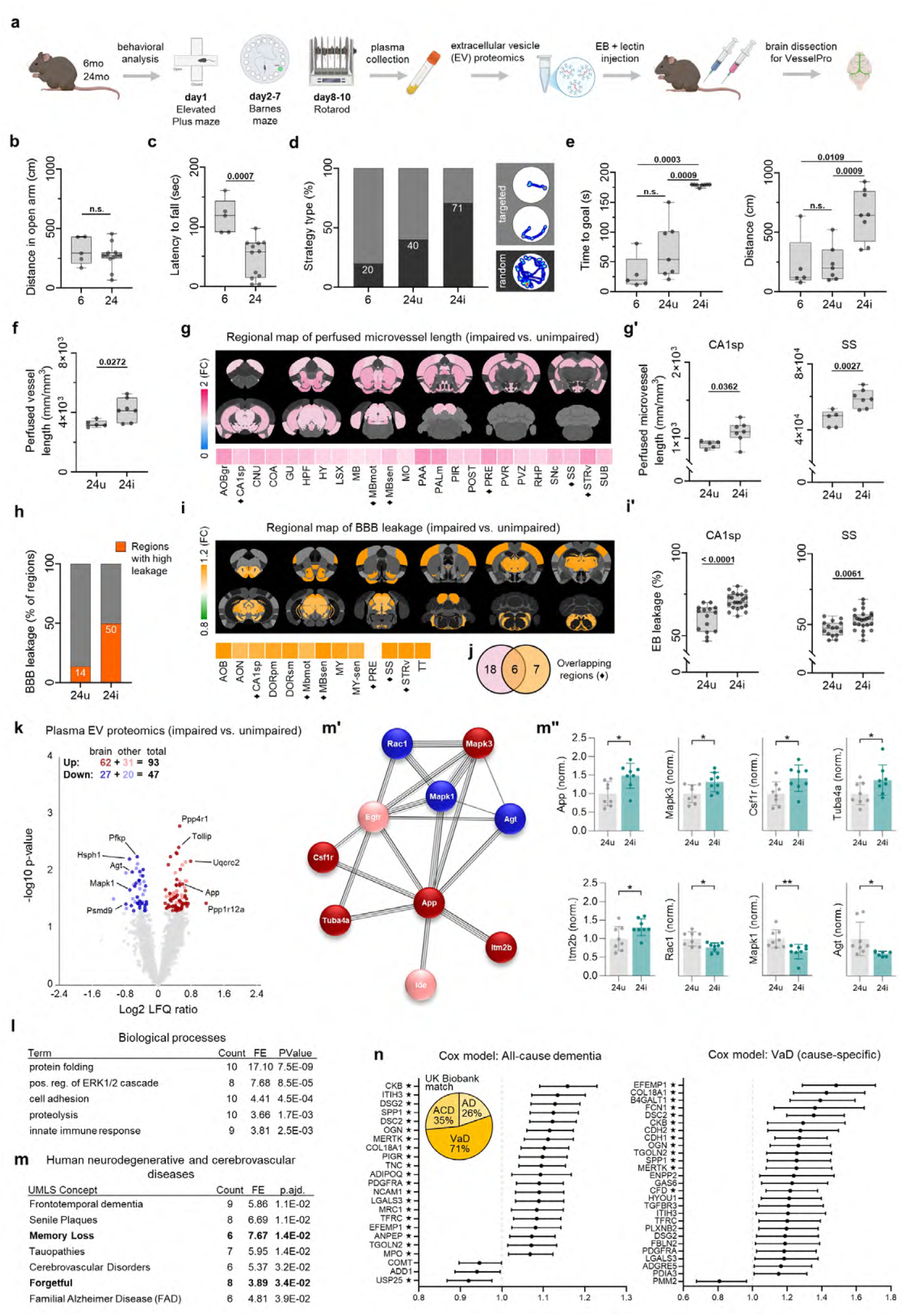
The hypervascular trajectory is associated with BBB leakage and spatial memory impairment. **a:** Schematic overview of the workflow used for the behavior testing (consisting of Elevated plus maze, Barnes maze (BM) and Rotarod tests) and the subsequent blood plasma collection and the VesselPro application on 6-months old and 24-months old mice. **b:** The Elevated plus maze test did not reveal a significant difference in the anxiety between the age groups. **c:** The Rotarod test showed that the 6-monthsold mice stayed longer on the rotating ramp. **d**,**e:** The distribution of the search strategies on the BM; the time spent and distance walked to find the goal split the aged cohort into cognitively impaired (24i) and unimpaired (24u) subgroups. The impaired subgroup showed a significantly slower goal search with longer distance traveled. **f:** Comparison of the perfused brain vessel length in the 24i and 24u mice. **g:** The 24i showed a consistent and statistically significant increase in perfused vessel length, primarily localized to cortical, striatal, hippocampal and mesoencephalic areas. **g’:** Quantifications of the perfused microvascular length in the CA1sp and the somatosensory cortex showing elevated vascular lengths in the 24i mice. **h-i’:** Quantification of the BBB leakage showing significant barrier failure in the 24i (**h**): primarily localized to cortical, hippocampal, thalamic and mesoencephalic areas (**i**) including the SS and the CA1sp (**i’**). **j:** Comparison of the brain regions with hypervascularization and/or leaky BBB reveals a partial overlap (common regions, marked with ♦ in g and i). **k:** Volcano plot of the blood plasma proteins in the 24i versus 24u. Proteins identified by the STRING DB as brain tissue expressed are shown in dark red and dark blue. **l:** The 5 most significant terms of the GO enrichment analysis of the altered proteins. **m-m”:** The 7 most significant human neurodegenerative and cerebrovascular diseases associated with the altered protein levels by the overrepresentation analysis (ORA). Protein-protein interaction network of the hits in the “Memory loss” and “Forgetful” concepts in STRING DB (node colors reflect the direction of abundance change as in k) and the quantification of the proteins with the most connections. The box and whisker plots show the median, data spread (whiskers), and individual data points; bar graphs show the group average ±SD (whiskers) and the individual data points. **n:** Distribution of shared plasma markers across All-cause dementia (ACD), Vascular dementia (VaD), and Alzheimer’s disease (AD) models (left). Intersection of significantly altered mouse plasma proteins with UK Biobank risk predictors for ACD (middle) and VaD (right). FDR significance marked with (★).

### Tie2 pathway activation reverses cognitive and barrier dysfunction and restores perfused vessel length

Based on the observed BBB permeability, the memory-impairment-associated hypervascularization, and elevated circulating vascular-inflammatory markers in aged, cognitively impaired mice, we tested whether pharmacological stabilization of the endothelium could restore vascular integrity and cognitive performance. We used AKB-9778, a small-molecule inhibitor of vascular endothelial protein tyrosine phosphatase (VE-PTP) that activates Tie2 signaling to enhance endothelial stability. Eighteen-month-old mice first completed a baseline Barnes maze (BM) test, then returned to their home cages for eight weeks. Animals were randomly assigned to AKB-9778 or vehicle treatment (4 days, twice daily s.c. injections). Following treatment, mice were re-trained and re-tested in the BM. Blood plasma was collected before treatment and at the final time point; brains were harvested for vascular and BBB analysis using VesselPro (**Fig. 5a**). AKB-9778 treated mice showed significant improvements in spatial memory compared with both baseline and vehicle-treated controls (**Fig. 5b**). Treated animals primarily used goal-directed (targeted) search strategies and had shorter latencies to find the escape hole (**Fig. 5c, Fig. S2b**). Vehicle-treated mice showed no change from baseline. VesselPro analysis showed that AKB-9778 treated animals displayed attenuated perfused vessel length relative to vehicle treated aged controls (**Fig. 5d**), resembling the vascular profile of cognitively unimpaired aged animals from the previous cohort (**Fig. 4f**). This effect, driven by the microvasculature (**Fig. 5e**) showed a heterogeneous pattern across 42 brain regions (**Fig. 5e’**). EB mapping revealed significantly improved BBB integrity post-treatment, with extravasation limited to 4.2% of brain regions, like in the 6-months-old mice and markedly lower than in the 6 months old brains, in stark contrast to the widespread leakage in untreated aged mice (**Fig. 5f**). Treatment with AKB-9778 resulted in reduced BBB leakage in 47 brain regions compared to the vehicle group (**Fig. 5f’**). Twenty-four brain regions showed both a reduced perfused vessel length and reduced BBB leakage (**Fig. 5g**). Interestingly, in several brain regions, we found that blood vessels displayed an increased proportion of open vessel segments (**Fig. 5h**), which could explain the decrease in microvascular length alongside the increased proportion of mesoscale vasculature. Nano-Mag-Net analysis of plasma extracellular vesicles revealed that AKB-9778 treatment reduced several brain-expressed markers, as indicated by STRING DB (**Fig. 5i**). This included the brain endothelium-specific proteins Flt4 (Vegfr-3) and Cdh5 (VE-cadherin), as well as brain mural cell-associated proteins Anpep, Pdgfrb, and Lama1 (**Fig. 5j**). Previous studies on the same mouse strain showed that AKB-9778 treatment increased these proteins in the neurovasculature (**Fig. 5k**, ^46^). Gene Set Enrichment Analysis (GSEA) revealed that the altered protein levels were associated with several memory and cerebrovascular-related diseases, including the “Memory Impairment” unified medical language system (UMLS) concept, involving 17 proteins (**Fig. 5l**). STRING DB analysis identified a cluster of 13 proteins forming a highly interactive network, with App and Tgf-β1 centrally positioned. All proteins in this cluster were down-regulated after AKB-9778 treatment, as reflected by the node colors (corresponding to the categories in the volcano plot) (**Fig. 5m**). Additionally, from the time-resolved proteomics data we found that markers associated with vessel growth and barrier integrity (Add1, Agt, Timp3, Tgfrb3, Sphk1, Thbs1, F11r, Gpld1, Itgb1) were elevated in the vehicle-treated group, consistent with pathological hypervascularization and increased brain permeability. In contrast, drug treatment attenuated their levels (**Fig. 5m’**). Together, these findings show that transient Tie2 activation is sufficient to attenuate the hypervascular and BBB-leaky phenotype and its associated cognitive deficits, consistent with endothelial instability acting as a key driver of this aging trajectory.

**Fig.5:**
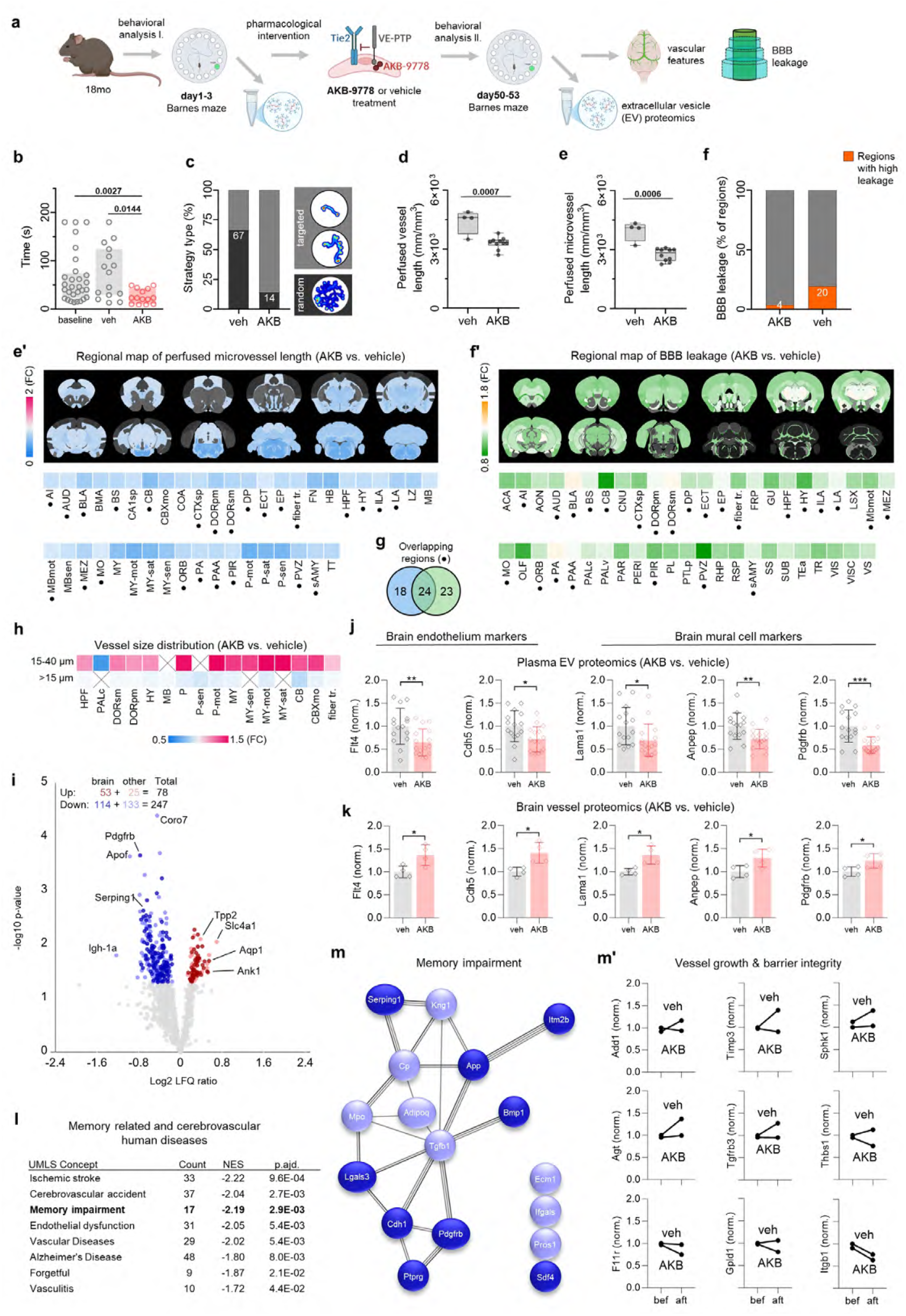
AKB-9778 treatment improves memory and BBB dysfunction restoring plasma proteomic signatures in aged mice. **a:** Schematic overview of the workflow used for the behavior testing (Elevated plus maze, Barnes maze (BM) and Rotarod tests) and the subsequent blood plasma collection and the VesselPro application on 18-months old mice. **b:** AKB-9778-treated aged mice exhibited significant improvement in spatial memory performance, compared to both baseline and vehicle-treated controls, by locating the escape hole consistently faster. **c:** Drug treated animals primarily employed goal-directed search strategies, whereas vehicle-treated mice showed no improvement. **d:** AKB-9778-treated mice had reduced perfused vessel length compared to vehicle-treated controls, resembling the vascular profile of cognitively unimpaired aged animals mainly contributed by (**e**) the microvessel length. **e’**: AKB treated mice showed a consistent and statistically significant decrease in perfused vessel length, primarily localized to hippocampal-, thalamic-, hypothalamic-, mesencephalic-, cerebellar- and to some cortical areas. **f:** BBB integrity was markedly improved following treatment limited to 4% of the brain regions, in contrast to the widespread leakage observed in untreated aged mice (20% of all brain regions). **f’:** The AKB treated mice showed a consistent and statistically significant decrease in BBB leakage, primarily localized to cortical-, hippocampal, thalam-ic-, and hypothalamic areas. **g:** Comparison of the brain regions with hypervascularization and/or leaky BBB reveals a partial overlap (24 common regions, marked with ● in e’ and f’). **h:** Comparison of the distribution of the perfused vessel sizes in the brain regions where the micro- and/or mesoscale vascular changes were apparent. The fold change of the significantly changed brain areas are colored on a red-blue scale, X marks non-significance. **i:** Volcano plot of the blood plasma proteins in the AKB-vs. vehicle-treated mice showing 78 significantly upregulated and 247 significantly downregulated proteins of which 53 and 114 respectively are identified by the STRING DB as brain tissue expressed (colored dark red and dark blue). **j:** Quantification of the dysregulated brain endothelial and mural cell marker proteins in the blood plasma of AKB-vs. vehicle-treated aged mice. **k:** Quantification of the dysregulated brain endothelial and mural cell marker proteins in the brain vessel proteomics AKB-vs. vehicle-treated adult mice from *Todorov-Völgyi et al*. **l:** The 8 most significant human memory related and cerebrovascular diseases associated with the altered protein levels by the gene set enrichment analysis (GSEA). **m:** Protein-protein interaction (ppi) network of the identified proteins of the “Memory impairment” concept in STRING DB. Node colors reflect the direction of abundance change as in (i). **m’:** Group average abundance levels of the vessel growth and barrier integrity related proteins before and after AKB or vehicle treatment, respectively. The box and whisker plots show the median, data spread (whiskers), and individual data points. The significantly changed brain areas are colored on a red-blue scale showing the fold change. Bar graphs show the group average ±SD (whiskers) and the individual data points.

## Discussion

Brain vascular aging does not follow a uniform decline: mice of the same chronological age segregate into distinct hypovascular and hypervascular trajectories with divergent functional outcomes. These trajectories are defined relative to a shared young-adult vascular baseline, thereby highlighting divergent vascular outcomes among aged individuals. This separation was supported across whole-brain vascular length metrics, regional mapping, and clustering analyses. The hypervascular phenotype identified here is not without precedent in humans, as regional hyperperfusion and vascular remodeling have been reported across vascular-driven brain pathologies, including vascular cognitive impairment and cerebral small vessel disease. Transient activation of the Tie2 pathway attenuated the hypervascular, BBB-leaky, and hyperperfusion-associated vascular state and improved cognitive performance, supporting endothelial instability as a causal driver of this aging trajectory. Previous whole-brain and region-specific imaging studies have reported that aging is associated, on average, with vascular rarefaction accompanied by increased vessel diameter and tortuosity. Our findings extend these observations by revealing substantial inter-individual heterogeneity that is masked by population-averaged analyses. While a subset of aged animals in our cohort exhibited the expected hypovascular phenotype consistent with vascular rarefaction, another subset displayed pronounced hypervascular remodeling characterized by increased perfused microvascular length and network complexity. Thus, vascular loss and dilation represent one prevalent outcome of cerebrovascular aging, whereas hypervascular remodeling constitutes an alternative phenotype associated with BBB dysfunction and cognitive impairment. At the molecular level, hypervascular mice showed enrichment of pathways related to angiogenesis, cytoskeletal remodeling, cell adhesion, and stress signaling, consistent with an activated and structurally unstable endothelium. At the molecular level, hypervascular mice showed enrichment of pathways related to angiogenesis, cytoskeletal remodeling, cell adhesion, and stress signaling, consistent with an activated and structurally unstable endothelium. Plasma EV proteomics mirrored these findings, revealing endothelial, mural, and inflammatory markers consistent with systemic vascular stress. While brain-wide structural hypervascularization has not been systematically quantified in human tissue, the plasma signature of the hypervascular, BBB-compromised state aligned strongly with the risk of vascular dementia in humans (UK Biobank), and showed substantially weaker concordance with Alzheimer’s disease signatures, identifying this phenotype as a VaD-like molecular endotype rather than a generic feature of neurodegeneration. Although our cross-sectional aging data do not resolve the temporal sequence of vascular remodeling and BBB dysfunction, the ability of acute Tie2 activation to simultaneously modulate vascular state, barrier integrity, and cognition constrains their causal relationship. It supports endothelial destabilization as a common upstream driver. Nevertheless, our study has limitations, including the use of a single mouse model, a cross-sectional design, and behavioral testing restricted to hippocampal-dependent memory; sex-specific effects were also not assessed. While the current implementation captures vascular state and BBB integrity at single time points, direct resolution of the temporal relationship between vascular remodeling and barrier dysfunction will require longitudinal extensions. Direct causal links for candidate pathways will require targeted perturbations. Future work should track vascular trajectories longitudinally and integrate human imaging and plasma profiling to validate and refine these findings. In conclusion, brain vascular aging follows discrete trajectories with distinct structural, molecular, and functional signatures. By integrating brain-wide vascular imaging, spatial proteomics, and endothelial perturbations, we identified a maladaptive hypervascular, BBB-leaky state that impairs cognition and can be acutely reversed by stabilizing the endothelium. These results provide a mechanistic framework for understanding heterogeneous vascular aging and highlight endothelial stability as a central regulator of neurovascular health in late life.

## Acknowledgements

This work was supported by the Vascular Dementia Research Foundation, Deutsche For-schungsgemeinschaft (DFG, German Research Foundation) under Germany’s Excellence Strategy within the framework of the Munich Cluster for Systems Neurology (EXC 2145 SyNergy, ID 390857198), CRC 1744 (ID 548585053); Leducq Foundation (22CVD01 to MD); ERA-NET Neuron (MatriSVDs, to MD), ERC Consolidator Grant (A.E., GA 865323), Nomis Heart Atlas Project Grant (A.E., Nomis Foundation). Some of the graphical illustrations used in the manuscript were prepared using BioRender.com. The authors used ChatGPT v4 (OpenAI) and Gemini v2.5 (Google) large language models to refine the language and style of this work, then carefully reviewed and edited the text accurately and took full responsibility for the final content.

## Author contributions

M.I.T. designed and performed most of the experiments and analyzed data; K.T.V., D.P.M., S.K., R.M., J.M., M.A., L.Z., A.N., J.C.P. performed experiments and analyzed data; L.L. contributed critical material for this study; H.S.B., M.S., M.K.G., M.D., F.H. and A.E. contributed critical input to study design and manuscript writing; A.E. initiated and coordinated the study, and wrote the manuscript together with M.I.T. and F.H.; all authors read and revised the manuscript.

## Materials and Methods

### Animals and housing

All experiments were conducted using male C57BL/6 mice obtained from Janvier Labs (France). Mice were aged in the animal facility of the Institute for Stroke and Dementia Research and housed in isolated ventilated HEPA filtered cages (12 h light/dark cycle with ad libitum access to food and water). Tissues were harvested in parallel and during the same daytime.

### Experimental design

The study was conducted in three main cohorts: Phenotyping cohort: 6-, 12-, and 24-month-old mice were used for the initial discovery of vascular phenotypes, supplemented by the few remaining survivors reaching 30 months of age. Brains were processed through the VesselPro pipeline for 3D vascular imaging, followed by spatial proteomic analysis. Cognitive cohort: 6- and 24-month-old mice underwent a battery of behavioral tests (Barnes maze, elevated plus maze, rotarod). Following behavior, plasma was collected, and brains were processed via the VesselPro pipeline to correlate vascular metrics and BBB integrity with cognitive performance. Intervention cohort: 18-month-old mice were first subjected to a baseline Barnes maze test after which plasma was collected. After an 8-week aging period, mice were randomly assigned to receive either the Tie2 activator AKB-9778 or a vehicle control. AKB-9778 (MedChem, Cat# HY-109041) was dissolved in sterile DMSO and Mygliol. Mice received twice-daily subcutaneous (s.c.) injections of either AKB-9778 at 10 mg/ kg or vehicle DMSO for 4 consecutive days. One day after the treatment, mice were retrained and re-tested on the Barnes maze, blood plasma was collected, and brains were processed via the VesselPro pipeline.

### Behavioral Assays

#### Barnes Maze Testing

The Barnes maze was used to assess learning and memory deficits in BL6 mice under two conditions: (i) at 24 vs. 6 months of age, and (ii) at 18 months of age following treatment with AKB-9778 or vehicle. Spatial memory was evaluated using a previously published shortened Barnes maze protocol with minor modifications^47^. Mice completed three phases of testing: habituation (days 1-2), training (day 2), and probe trials (day 3) for short-term memory. All mice were acclimated to the testing room for 1 h prior to each session. During habituation, mice were placed in the center of the maze under a glass beaker for 30 s. They were then slowly guided to the target hole leading to the escape cage by moving the beaker over 10-15 s. Once positioned at the target hole, mice were given 1 min to enter the escape cage independently. If they did not enter on their own, they were gently nudged with the beaker. After entering, mice remained in the escape cage for 1 min before being returned to the holding cage. Each mouse completed three habituation sessions in total (two on the afternoon of day 1 and one on the morning of day 2). For training, mice were again placed in the center of the maze under a glass beaker for 30 s. After this period the beaker was removed, and mice were allowed to explore the maze for up to 3 min. If a mouse located the target hole and entered the escape cage, it was allowed to remain there for 1 min before being returned to the holding cage. If it failed to find the escape hole, it was gently guided toward it using the beaker. Should the mouse not enter the escape cage within 1 min of guidance, it was nudged with the beaker; if it still did not enter, it was manually placed into the escape cage and allowed to remain for 1 min before being returned to the holding cage. For probe testing, mice were placed in the center of the maze under a glass beaker for 30 s and then given 3 min to explore the maze. At the end of the trial, mice were returned to their holding cages. Latency and path length to reach the target hole were recorded and quantified as the primary measure of spatial memory.

#### Rotarod

Motor coordination and balance were assessed using the Rotarod test. Mice were trained over two consecutive days prior to performance testing. On day 1, mice completed two trials at a constant low speed (5 rpm) followed by three trials on an accelerating protocol (5-30 rpm over 300 s). On day 2, mice underwent three additional training trials using the same accelerating protocol (5-30 rpm over 300 s). These sessions served to familiarize animals with the apparatus and reduce variability due to novelty or anxiety. On day 3, motor performance was evaluated across three trials using the accelerating speed protocol (5-30 rpm over 300 s). For each trial, latency to fall from the rotating rod was recorded as the primary outcome measure. For each mouse, the mean latency across the three testing trials was calculated and used for statistical analysis.

#### Elevated plus maze

Anxiety-like behavior was assessed using the Elevated Plus Maze (EPM), a cross-shaped apparatus consisting of two open- and two closed arms elevated 50-60 cm above the floor. Prior to testing, mice were acclimated to the behavioral testing room for 1 h. During the testing procedure each mouse was placed in the center of the maze under a glass beaker for 30 s. Next, mice were allowed to freely explore the maze for 5 min. The time spent in open versus closed arms was recorded and quantified as the primary measure of anxiety-like behavior.

### VesselPro Pipeline: Perfusion, Clearing, and Imaging

#### Fluorescent labeling of the perfused cerebrovasculature and BBB leakage

To assess BBB permeability, mice were injected intraperitoneally (i.p.) with Evans Blue (EB) dye (2% in saline, 4 ml/kg) 12 h prior to sacrifice. Mice were deeply anesthetized using midazolam + medetomidine + fentanyl (i.p.; 1 ml per 100 g body mass for mice; 5 mg, 0.5 mg and 0.05 mg per kg body mass) and their thorax opened and transcardially injected with a solution containing wheat germ agglutinin (WGA) conjugated to AlexaFluor 594 (Thermo Fisher Scientific, W11262) 0.25 mg/ml in 150 μl PBS (pH 7.2) to label the functional, perfused vasculature. After cervical dislocation the brains were harvested and placed into a 5 ml tube with 4% PFA for 6 h.

#### Optical tissue clearing

Brains were optically cleared using an adapted 3DISCO protocol^48^. Briefly, we immersed them in a gradient of tetrahydrofuran (Sigma-Aldrich, 186562): 50 vol%, 70 vol%, 80 vol%, 90 vol%, 100 vol% (in distilled water), and 100 vol% at room temperature for 12 h at each concentration, delipidated in dichloromethane (Sigma-Aldrich, 270997) for 12 h at room temperature and finally incubation with the refractive index matching solution BABB (benzyl alcohol + benzyl benzoate 1:2 ratio; Sigma-Aldrich, 24122 and W213802), for at least 24 h at room temperature until transparency was achieved. Each incubation step was carried out on a laboratory shaker.

#### Light-sheet fluorescence microscopy

We used a 4× objective lens (Olympus XLFLUOR 340) equipped with an immersion corrected dipping cap mounted on a LaVision UltraII microscope coupled to a white light laser module (NKT SuperK Extreme EXW-12) for imaging. The images were taken in 16 bit depth and at a nominal resolution of 1.625 μm / voxel on the XY axes. The brain vasculature was visualized by the Alexa Fluor 594 (using a 580/25 nm excitation and a 625/30 nm emission filter) and Evans blue fluorescent dyes (using a 640/40 nm excitation and a 690/50 nm emission filter) in a sequential order. We maximized the signal to noise ratio (SNR) for each dye independently to avoid saturation of differently sized vessels when only a single channel is used. We achieved this by independently optimizing the excitation power so that the strongest signal in the major vessels does not exceed the dynamic range of the camera. To reduce defocus, which derives from the Gaussian shape of the beam, we used a 12-step sequential shifting of the focal position of the light-sheet per plane and side. Scans were done using grid-tiling with 20% overlap in both X- and Y-directions to encompass the whole tissue. In z-dimension we took the sectional images in 3 μm steps using left and right sided illumination.

### Image Processing and Quantification

#### 3D vascular analysis

We used TeraStitcher’s automatic global optimization function (v1.10.10) for 3D data reconstruction from the tiling volumes. To register our dataset to the reference atlas we used the average template, the annotation file, and the ontology description of the Allen Mouse Brain Common Coordinate Framework (CCFv3)^49^. Brain vasculature was detected and quantified using our previously established Vessel Segmentation and Analysis Pipeline^43^ (VesSAP). Quantitative metrics included total vessel length normalized to whole-brain volume, regional vessel length normalized to brain region volume, bifurcation density (number of branch points normalized to region volume), and the radius of the vessels. All measurements were corrected by a constant factor to account for tissue shrinkage introduced by fixation and clearing. 3D renderings were made with Imaris 9.9 (Oxford Instruments).

#### Vascular loop detection

We based our analysis on the previously published and established vessel graph extraction and analysis pipeline ^50^, which combines graph extraction ^51^ with a subsequent post-processing step for graph pruning and feature merging. To assess and statistically quantify vessel topology, we extend the pipeline by implementing further processing steps. For this, we cast the extracted graph to an undirected graph representation using the PyTorch Geometrics^52^ and NetworkX ^53^ frameworks. In this undirected graph, nodes correspond to vessel bifurcations while edges constitute vessel segments. First, we preprocess the graph by pruning vessel segments, filtering segments shorter than 5 voxels, and subsequently removing isolated nodes and self-loops. Finally, we identify vascular loops by computing the cycle basis of the graph, i.e., the minimal set of cycles that form a basis for the cycle space, by employing the implementation in NetworkX. We then filter the detected cycles to retain only loops consisting of between 3 and 15 vessel segments (edges).

#### BBB leakage quantification

Leakage analysis was done by generating three-dimensional concentric shells around the vessel segmentation mask, with shell distances of 2, 4, and 6 voxels, respectively. The mean intensity of the EB channel was computed for the voxels within each shell (excluding those with zero intensity values) and normalized to the mean intensity of the EB in the vascular mask (denoted as the shell at voxel 0) in the major brain regions. The BBB leakage of the major brain regions was averaged between hemispheres.

### Proteomic Analysis

#### Spatial brain tissue proteomics and data analysis

Following imaging, cleared brains were rehydrated through a reverse-graded tetrahydrofuran series, embedded in 4% agarose and sliced with a vibratome (Leica, Germany) into 100 μm thick coronal sections. Brain regions (hippocampus, somatosensory cortex) were anatomically sampled under an Axio Zoom microscope (Zeiss, Germany) based on the 3D imaging results with a Sterican 18G stainless steel syringe needle (Braun, Germany). Sample preparation for proteomics analysis was performed as described previously with slight modifications ^44^. Briefly, the samples were first resuspended in 6% SDS buffer, heat denatured at 95°C for 45 min at 600 rpm in a thermoshaker, sonicated in high mode for 30 cycles (30 s OFF, 30 s ON) (Bioruptor Plus, Diagenode) and then precipitated using 80% acetone overnight in -20°C. The next day, these samples were centrifuged and the pellet was resuspended in SDC lysis buffer (2% SDC, 100 mM Tris-HCl pH 8.5). The samples in the SDC buffer were sonicated in high mode for 15 cycles (30 s OFF, 30 s ON). The samples were again heated at 95°C for 45 min at 600 rpm in a thermoshaker. The protein samples were digested overnight with Trypsin and LysC (1:50, protease:protein ratio) at 37°C, 1,000 rpm shake. Resulting peptides were acidified with 1% TFA 99% isopropanol with 1:1 volume-to-volume ratio, vortexed and centrifuged to pellet residual particles. The supernatant was transferred to fresh tubes and subjected to in-house built StageTip clean-up consisting of three layers of styrene divinylbenzene reversed-phase sulfonate (SDB-RPS; Empore, 3M) membranes. Peptides were loaded on the activated (100% ACN, 1% TFA in 30% MeOH, 0.2% TFA, respectively) StageTips, run through the SDB-RPS membranes, and washed by EtOAc including 1% TFA, isopropanol including 1% TFA, and 0.2% TFA, respectively. Peptides were then eluted from the membranes via 60 µL elution buffer (80% ACN, 1.25% NH4OH) and dried using a vacuum centrifuge (40 min at 45°C). Finally, peptides were reconstituted in 8-10 µl of loading buffer (2% ACN, 0.1% TFA) and stored in -80 until further use. The mass spectrometry data was acquired in data-independent acquisition (DIA) mode. The LC-MS/MS analysis was carried out using EASY nanoLC 1200 (Thermo Fisher Scientific) coupled with trapped ion mobility spectrometry quadrupole time-of-flight single cell proteomics mass spectrometer (timsTOF SCP, Bruker Daltonik GmbH, Germany) via a CaptiveSpray nano-electrospray ion source. Peptides (50 ng) were loaded onto a 25 cm Aurora Series UHPLC column with CaptiveSpray insert (75 μm ID, 1.6 μm C18) at 50°C and separated using a 50 min gradient (5-20% buffer B in 30 min, 20-29% buffer B in 9 min, 29-45% in 6 min, 45-95% in 5 min, wash with 95% buffer B for 5 min, 95-5% buffer B in 5 min) at a flow rate of 300 nL/min. Buffer A and B were water with 0.1 vol% formic acid and 80:20:0.1 vol% ACN:water: formic acid, respectively. MS data were acquired in single-shot library-free DIA mode and the timsTOF SCP was operated in DIA/parallel accumulation serial fragmentation (PASEF) using the high sensitivity detection-low sample amount mode. The ion accumulation and ramp time were set to 100 ms each to achieve nearly 100% duty cycle. The collision energy was ramped linearly as a function of the mobility from 59 eV at 1/K0 = 1.6 Vs cm^−2^ to 20 eV at 1/K0 = 0.6 Vs cm^−2^. The isolation windows were defined as 24 × 25 Th from m/z 400 to 1000. diaPASEF raw files were searched against the mouse UniProt database using DIA-NN ^54^. Peptides with a length of at least seven amino acids were considered for the search including N-terminal acetylation. Oxidation of methionine was set as a variable modification and cysteine carbamidomethylation as fixed modification. Enzyme specificity was set to Trypsin/P with 2 missed cleavages. The FASTA digest for library-free search was enabled for predicting the library generation. The FDR was set to 1% at precursor and global protein level. Match-between-runs (MBR) feature was enabled and quantification mode was set to “Robust LC (high precision)”. The Protein Group column in DIA-NN’s report was used to identify the protein group and PG.Max-LFQ was used to calculate the differential expression.

### Plasma derived extracellular vesicle (EV) proteomics

#### Mouse plasma preparation

From the cognitive and intervention animal cohorts blood samples were collected from the tail vein of the mice into blood collection tubes containing EDTA anticoagulant (Sarstedt). Plasma was separated from the blood cells by centrifugation with 2000 x g for 10 min at 4 °C and then stored at -80 °C until further processing. Nano-Mag-Net capture of plasma particles Negatively charged particles contained were enriched from mouse plasma using a simplified Mag-Net^45^ protocol. Briefly, buffer exchanged strong anion exchange (SAX) beads were prepared in a Nano-Mag-Net (NMN) buffer containing 50% (v/v) Phosphate Buffered Saline and Tris-HCl (pH 8.5, 250 mM). Mouse plasma (2.5 μl) was incubated with 0.5 μl of these NMN buffer pre-equilibrated SAX beads for 15 minutes to allow capture of negatively charged particles. The plasma particle-loaded SAX beads were then washed four times with 20 μl of NMN buffer. Beads were captured on a magnetic 96-well plate, with supernatants discarded between washes. The beads were mixed by shaking and spun down for recovery. Subsequently, the beads underwent reduction, alkylation, and denaturation (RAD) for 10 min at 90 °C on a heat block in RAD buffer containing 2% SDC, 10 mM TCEP, 40 mM CAA, and 100 mM Tris-HCl. Protein cleanup was performed using SP3^55^ by precipitating proteins in 30 μl ethanol for 5 min, followed by three rounds of washing in 20 μl of 80% ethanol. The precipitated proteins were dried in a SpeedVac at 60 °C. Finally, proteins were digested using 500 ng Trypsin-LysC in 25 μl of a digestion mix containing 50 mM Hepes (pH 7.5), 1 mM CaCl2, and 0.015% DDM.

#### Evotip PURE clean-up of plasma particle samples

Miniaturised, 96-well-scalable C18 cleanup was performed as described previously^56^. Briefly, Evotip PURE tips were rinsed with 20 μl of Buffer B (comprising 80% ACN, water and 0.1% formic acid) and spun down at 800 x g for 60 s. The Evotips were conditioned with 10 μl of isopropanol, followed by an impulse spin at 100 x g, a 1-min incubation and an additional 4 min at 100 x g to empty the Evotips. The PURE Evotips were equilibrated in 20 μl of Buffer A (0.1% formic acid) and impulse spun at 800 x g for storage until the acidified samples were ready to load. Plasma particle samples were acidified in 1% TFA, and the Evotip PURE was emptied by centrifuging at 800 x g for 1 min. The acidified samples were loaded onto the PURE Evotips and spun at 800 x g for 1 min. The samples were washed twice with 20 μl of Buffer A and spun down at 800 x g for 1 min. Elutions were collected in PCR strips by eluting with 20 μl of 45% Buffer B (containing 45% ACN, water and 0.1% TFA) at 450 x g. The peptides were dried in a SpeedVac and resuspended in 0.1% TFA supplemented with 0.015% DDM for mass spectrometry analyses on the timstofSCP platform as described previously^56^ using a 5.5 cm uPAC HT column (Thermo Scientific) on the EASY nLC chromatography coupled to a Bruker timstofSCP using diaPASEF.

#### Proteomics Data Preprocessing

Raw proteomic intensities were subjected to a rigorous preprocessing pipeline using scanpy^57^ to ensure data quality and statistical robustness. To minimize the impact of stochastic detection, we first applied a stringent filtering criterion, retaining only those proteins consistently identified in at least 5 samples per experimental group. Next, the data were normalized per sample to account for variations in total protein loading and instrument sensitivity. The normalized intensity values were subsequently log1p-transformed (log plus one) to achieve a normal distribution and stabilize variance. To address the remaining missing values inherent in bottom-up proteomics, we employed k-Nearest Neighbors (kNN) imputation, which utilizes the local similarity between protein profiles to provide biologically informed estimates for missing data points. This processed dataset served as the foundation for all subsequent differential expression and pathway enrichment analyses. Enrichment analysis of biological processes (GOTERM_BP_DIRECT) was performed with Database for Annotation, Visualization and Integrated Discovery (DAVID) tool^58^, and protein-protein-interactions and tissue expression analyses were done with the STRING DB (Search Tool for Recurring Instances of Neighbouring Genes, v12.0) tool^59^. In both tools the Mus musculus standard background dataset was used.

### Disease Association Analysis

Differentially abundant proteins were converted to log2 fold-changes, and used to generate per-experiment ranked protein lists. Where indicated, mouse symbols were mapped to high-confidence human orthologs using gprofiler^60^ removing unmapped entries and resolving duplicate human mappings by retaining the gene with the largest absolute effect size. Enrichment was performed adaptively: datasets with a sufficiently large fold-change range were analyzed by gene set enrichment analysis (GSEA) using clusterProfiler^61^ for Gene Ontology (GO) Biological Processes, KEGG pathways, and DisGeNET diseases, applying Benjamini-Hochberg multiple-testing correction. For datasets with small effect sizes or extensive ties, over-representation analysis (ORA) was used instead, testing up- and down-regulated gene sets defined by log_2_FC thresholds against the experiment-specific background. All analyses were performed in R (v4.5.1) using the following packages and versions: tidyverse (v2.0.0), here (v1.0.2), fs (v1.6.6), glue (v1.8.0), clusterProfiler (v4.16.0), enrichplot (v1.28.4), DOSE (v4.2.0), org.Mm.eg.db (v3.21.0), org.Hs.eg.db (v3.21.0), and gprofiler2 (v0.2.4).

### UK Biobank analysis

#### Human cohort and proteomics

We utilized Olink proteomic data from the UK Biobank (UKB) Pharma Proteomics Project. In total, up to 2,923 proteins were measured in EDTA plasma samples from approximately 54,219 participants using the Olink Explore 3072 platform. Protein abundances are reported as Normalized Protein eXpression (NPX) values on a log_2_ scale.

#### Statistical model

We used Cox proportional hazards regression to evaluate associations between each protein and three dementia outcomes ascertained from UKB algorithmically defined outcomes: all-cause dementia (ACD; Field 42018), vascular dementia (VaD; Field 42022), and Alzheimer’s disease (AD; Field 42020). Models were adjusted for baseline age and sex, and participants were followed for dementia diagnoses through October 31 2022. Missing values were not imputed, and each protein was analyzed using complete cases only.

#### Cross-species comparison

We compared the results of our mouse plasma proteomics (Vehicle-treated vs. AKB-9778-treated) with the human dementia Cox models. Mouse proteins were first mapped to human orthologs using UniProt identifiers; only proteins quantified on both platforms were included in the comparison. A protein was considered a “match” if it was significantly associated with disease risk in the human data (p < 0.05). We then assessed “concordant directionality,” defined as the mouse fold-change (FC) and the human hazard ratio (HR) both being > 1 or both being < 1.

### Statistical analysis

Unless otherwise specified, statistical analyses were performed using GraphPad Prism (v10.5) or Python’s (v3.8) scipy.stats package (v1.10.0). Unsupervised clustering methods (MiniBatch KMeans, Ward, Agglomerative Clustering, OPTICS, BIRCH, Gaussian Mixture) were used from the scikit-learn (v1.1.2) package. Comparisons between two groups were made using unpaired Student’s t-tests with Welch’s correction. Correlations were assessed using Pearson’s or robust linear regression. Data are presented as mean ± SEM, and significance was set at p < 0.05.

---

**Fig. S1.**
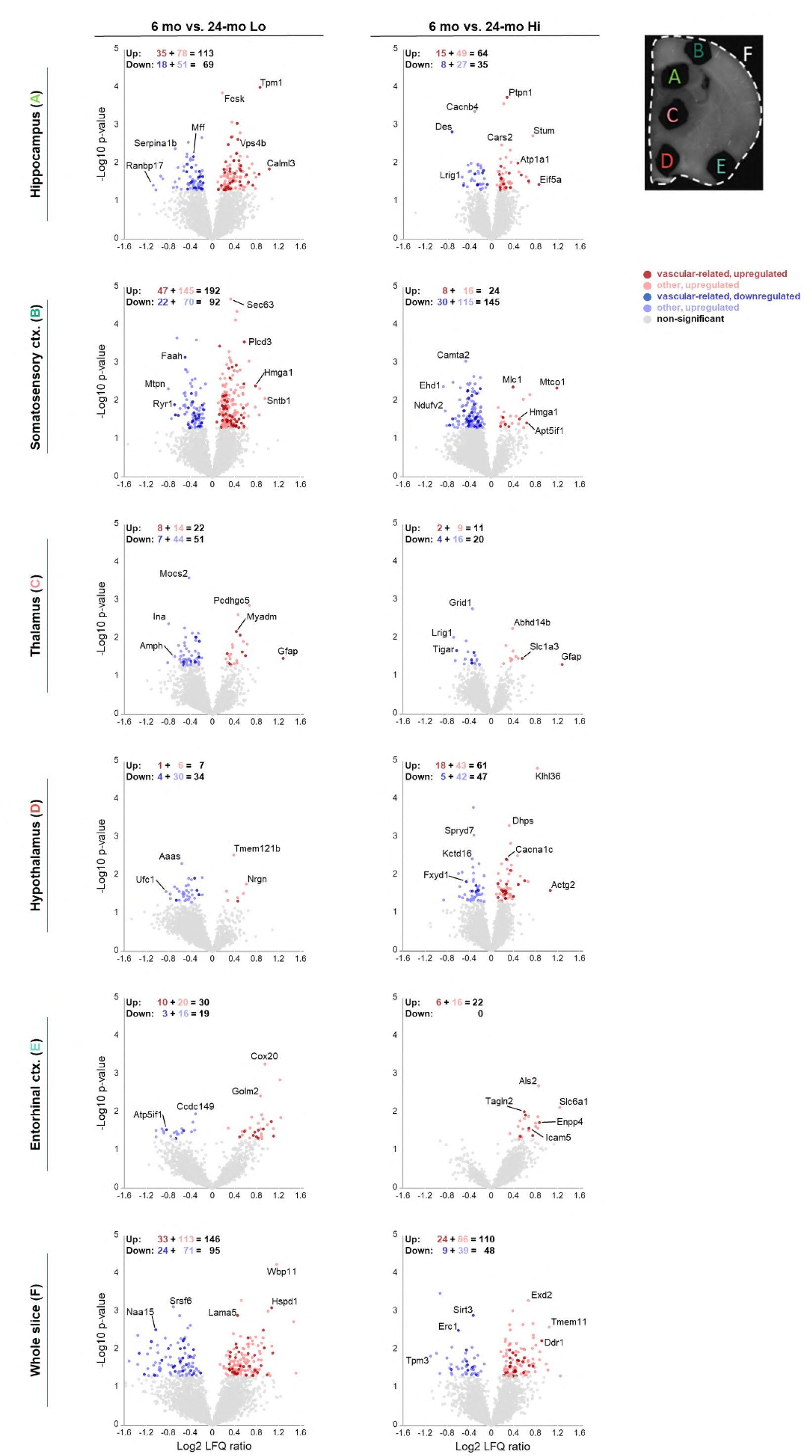
Volcano plots and quantification of the differentially expressed proteins across different brain regions. Vascular-related proteins are shown with dark red and blue.

**Fig. S2.**
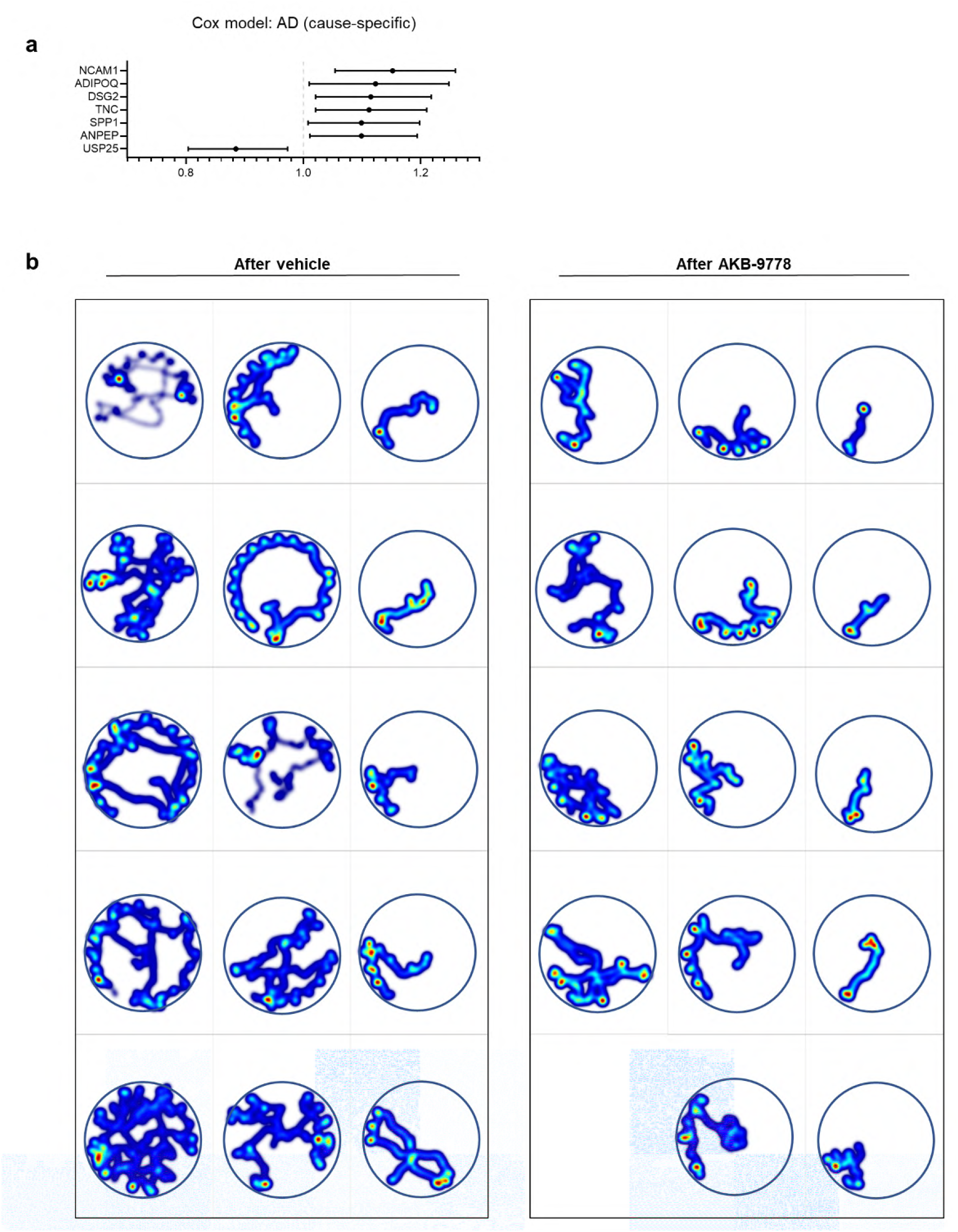
**a:** Intersection of significantly altered mouse plasma proteins and significant predictors of AD risk in the UK Biobank (cause-specific model). **b:** Heatmaps of the test traces for individual mice after AKB-9778 or vehicle treatment.

## References

1. Iadecola, C. The pathobiology of vascular dementia. Neuron 80, 844–866 (2013).

2. Wardlaw, J. M., Smith, C. & Dichgans, M. Small vessel disease: mechanisms and clinical implications. Lancet Neurol. 18, 684–696 (2019).

3. Wardlaw, J. M., Sandercock, P. a. G., Dennis, M. S. & Starr, J. Is breakdown of the blood-brain barrier responsible for lacunar stroke, leukoaraiosis, and dementia? Stroke 34, 806–812 (2003).

4. Sweeney, M. D., Zhao, Z., Montagne, A., Nelson, A. R. & Zlokovic, B. V. Blood-Brain Barrier: From Physiology to Disease and Back. Physiol. Rev. 99, 21–78 (2019).

5. Taheri, S. et al. Blood-brain barrier permeability abnormalities in vascular cognitive impairment. Stroke 42, 2158–2163 (2011).

6. Zhang, C. E. et al. Blood-brain barrier leakage in relation to white matter hyperintensity volume and cognition in small vessel disease and normal aging. Brain Imaging Behav. 13, 389–395 (2019).

7. Montagne, A. et al. Blood-brain barrier breakdown in the aging human hippocampus. Neuron 85, 296–302 (2015).

8. Sweeney, M. D., Sagare, A. P. & Zlokovic, B. V. Blood-brain barrier breakdown in Alzheimer disease and other neurodegenerative disorders. Nat. Rev. Neurol. 14, 133– 150 (2018).

9. Nation, D. A. et al. Blood–brain barrier breakdown is an early biomarker of human cognitive dysfunction. Nat. Med. 25, 270–276 (2019).

10. Alsop, D. C., Casement, M., de Bazelaire, C., Fong, T. & Press, D. Z. Hippocampal Hyperperfusion in Alzheimer’s Disease. NeuroImage 42, 1267–1274 (2008).

11. Thomas, K. R. et al. Regional hyperperfusion in older adults with objectively-defined subtle cognitive decline. J. Cereb. Blood Flow Metab. 41, 1001–1012 (2021).

12. Ding, B. et al. Pattern of cerebral hyperperfusion in Alzheimer’s disease and amnestic mild cognitive impairment using voxel-based analysis of 3D arterial spin-labeling imaging: initial experience. Clin. Interv. Aging 9, 493–500 (2014).

13. Callewaert, B., Gsell, W., Lox, M., Himmelreich, U. & Jones, E. A. V. A timeline study on vascular co-morbidity induced cerebral endothelial dysfunction assessed by perfusion MRI. Neurobiol. Dis. 202, 106709 (2024).

14. Wang, X. et al. Brain hemodynamic changes in amnestic mild cognitive impairment measured by pulsed arterial spin labeling. Aging 12, 4348–4356 (2020).

15. Wierenga, C. E. et al. Effect of mild cognitive impairment and APOE genotype on resting cerebral blood flow and its association with cognition. J. Cereb. Blood Flow Metab. 32, 1589–1599 (2012).

16. Tong, X.-K., Nicolakakis, N., Kocharyan, A. & Hamel, E. Vascular Remodeling versus Amyloid β-Induced Oxidative Stress in the Cerebrovascular Dysfunctions Associated with Alzheimer’s Disease. J. Neurosci. 25, 11165–11174 (2005).

17. Liu, X. et al. Progressive mechanical and structural changes in anterior cerebral arteries with Alzheimer’s disease. Alzheimers Res. Ther. 15, 185 (2023).

18. Shang, J. et al. Strong Impact of Chronic Cerebral Hypoperfusion on Neurovascular Unit, Cerebrovascular Remodeling, and Neurovascular Trophic Coupling in Alzheimer’s Disease Model Mouse. J. Alzheimers Dis. JAD 52, 113–126 (2016).

19. Zhang, P. et al. Early exercise improves cerebral blood flow through increased angiogenesis in experimental stroke rat model. J. NeuroEngineering Rehabil. 10, 43 (2013).

20. Rust, R. et al. Anti-Nogo-A antibodies prevent vascular leakage and act as pro-angiogenic factors following stroke. Sci. Rep. 9, 20040 (2019).

21. Desai, B. S., Schneider, J. A., Li, J.-L., Carvey, P. M. & Hendey, B. Evidence of angiogenic vessels in Alzheimer’s disease. J. Neural Transm. Vienna Austria 1996 116, 587– 597 (2009).

22. Burke, M. J. C. et al. Morphometry of the hippocampal microvasculature in post-stroke and age-related dementias. Neuropathol. Appl. Neurobiol. 40, 284–295 (2014).

23. Grammas, P. Neurovascular dysfunction, inflammation and endothelial activation: Implications for the pathogenesis of Alzheimer’s disease. J. Neuroinflammation 8, 26 (2011).

24. Carmeliet, P. & Ruiz de Almodovar, C. VEGF ligands and receptors: implications in neurodevelopment and neurodegeneration. Cell. Mol. Life Sci. CMLS 70, 1763– 1778 (2013).

25. Shim, J. W. & Madsen, J. R. VEGF Signaling in Neurological Disorders. Int. J. Mol. Sci. 19, 275 (2018).

26. Tarkowski, E. et al. Increased intrathecal levels of the angiogenic factors VEGF and TGF-beta in Alzheimer’s disease and vascular dementia. Neurobiol. Aging 23, 237– 243 (2002).

27. Zhang, Z. G. et al. VEGF enhances angiogenesis and promotes blood-brain barrier leakage in the ischemic brain. J. Clin. Invest. 106, 829–838 (2000).

28. Hu, Y., Zheng, Y., Wang, T., Jiao, L. & Luo, Y. VEGF, a Key Factor for Blood Brain Barrier Injury After Cerebral Ischemic Stroke. Aging Dis. 13, 647–654 (2022).

29. Wu, M. et al. VEGF regulates the blood-brain barrier through MMP-9 in a rat model of traumatic brain injury. Exp. Ther. Med. 24, 728 (2022).

30. Lochhead, J. J., Yang, J., Ronaldson, P. T. & Davis, T. P. Structure, Function, and Regulation of the Blood-Brain Barrier Tight Junction in Central Nervous System Disorders. Front. Physiol. 11, 914 (2020).

31. Luissint, A.-C., Artus, C., Glacial, F., Ganeshamoorthy, K. & Couraud, P.-O. Tight junctions at the blood brain barrier: physiological architecture and disease-associated dysregulation. Fluids Barriers CNS 9, 23 (2012).

32. Dithmer, S., Blasig, I. E., Fraser, P. A., Qin, Z. & Haseloff, R. F. The Basic Requirement of Tight Junction Proteins in Blood-Brain Barrier Function and Their Role in Pathologies. Int. J. Mol. Sci. 25, 5601 (2024).

33. Antonetti, D. A., Barber, A. J., Hollinger, L. A., Wolpert, E. B. & Gardner, T. W. Vascular endothelial growth factor induces rapid phosphorylation of tight junction proteins occludin and zonula occluden 1. A potential mechanism for vascular permeability in diabetic retinopathy and tumors. J. Biol. Chem. 274, 23463–23467 (1999).

34. Bates, D. O. Vascular endothelial growth factors and vascular permeability. Cardiovasc. Res. 87, 262–271 (2010).

35. Yan, F. et al. Dimension-based Quantification of Aging-Associated Cerebral Microvasculature Determined by Optical Coherence Tomography and Two-Photon Microscopy. J. Biophotonics 17, e202300409 (2024).

36. Quintana, D. D. et al. The Cerebral Angiome: High Resolution MicroCT Imaging of the Whole Brain Cerebrovasculature in Female and Male Mice. NeuroImage 202, 116109 (2019).

37. Liebmann, T. et al. Three-dimensional study of Alzheimer’s disease hallmarks using the iDISCO clearing method. Cell Rep. 16, 1138–1152 (2016).

38. Di Giovanna, A. P. et al. Whole-brain vasculature reconstruction at the single capillary level. Sci. Rep. 8, 1–11 (2018).

39. d’Esposito, A. et al. Computational fluid dynamics with imaging of cleared tissue and of in vivo perfusion predicts drug uptake and treatment responses in tumours. Nat. Biomed. Eng. 2, 773–787 (2018).

40. Bennett, H. C. et al. Aging drives cerebrovascular network remodeling and functional changes in the mouse brain. Nat. Commun. 15, 6398 (2024).

41. Kirst, C. et al. Mapping the Fine-Scale Organization and Plasticity of the Brain Vasculature. Cell 180, 780-795.e25 (2020).

42. Shi, S. M. et al. Glycocalyx dysregulation impairs blood-brain barrier in ageing and disease. Nature 639, 985–994 (2025).

43. Todorov, M. I. et al. Machine learning analysis of whole mouse brain vasculature. Nat. Methods 17, 442–449 (2020).

44. Bhatia, H. S. et al. Spatial proteomics in three-dimensional intact specimens. Cell 185, 5040-5058.e19 (2022).

45. Wu, C. C. et al. Enrichment of extracellular vesicles using Mag-Net for the analysis of the plasma proteome. Nat. Commun. 16, 5447 (2025).

46. Todorov-Völgyi, K. et al. The stroke risk gene Foxf2 maintains brain endothelial cell function via Tie2 signaling. Nat. Neurosci. https://doi.org/10.1038/s41593-025-02136-5 (2025) doi:10.1038/s41593-025-0213

47. Attar, A. et al. A Shortened Barnes Maze Protocol Reveals Memory Deficits at 4-Months of Age in the Triple-Transgenic Mouse Model of Alzheimer’s Disease. PLoS ONE 8, e80355 (2013).

48. Ertürk, A. et al. Three-dimensional imaging of solvent-cleared organs using 3DISCO. Nat. Protoc. 7, 1983–1995 (2012).

49. Wang, Q. et al. The Allen Mouse Brain Common Coordinate Framework: A 3D Reference Atlas. Cell 181, 936-953.e20

50. Paetzold, J. C. et al. Whole Brain Vessel Graphs: A Dataset and Benchmark for Graph Learning and Neuroscience (VesselGraph). 10.5281/zenodo.5301621 (2021).

51. Drees, D., Scherzinger, A., Hägerling, R., Kiefer, F. & Jiang, X. Scalable Robust Graph and Feature Extraction for Arbitrary Vessel Networks in Large Volumetric Datasets. BMC Bioinformatics 22, 346 (2021).

52. Fey, M. & Lenssen, J. E. Fast Graph Representation Learning with PyTorch Geometric. Preprint at 10.48550/arXiv.1903.02428 (2019).

53. Hagberg, A. A., Schult, D. A. & Swart, P. J. Exploring Network Structure, Dynamics, and Function using NetworkX. in Proceedings of the 7th Python in Science Conference (eds Varoquaux, G., Vaught, T. & Millman, J.) 11–15 (Pasadena, CA USA, 2008).

54. Demichev, V., Messner, C. B., Vernardis, S. I., Lilley, K. S. & Ralser, M. DIA-NN: neural networks and interference correction enable deep proteome coverage in high throughput. Nat. Methods 17, 41–44 (2020).

55. Hughes, C. S. et al. Single-pot, solid-phase-enhanced sample preparation for proteomics experiments. Nat. Protoc. 14, 68–85 (2019).

56. Luo, J. et al. Nanocarrier imaging at single-cell resolution across entire mouse bodies with deep learning. Nat. Biotechnol. 43, 2009–2022 (2025).

57. Wolf, F. A., Angerer, P. & Theis, F. J. SCANPY: large-scale single-cell gene expression data analysis. Genome Biol. 19, 15 (2018).

58. Huang, D. W., Sherman, B. T. & Lempicki, R. A. Systematic and integrative analysis of large gene lists using DAVID bioinformatics resources. Nat. Protoc. 4, 44–57 (2009).

59. Snel, B., Lehmann, G., Bork, P. & Huynen, M. A. STRING: a web-server to retrieve and display the repeatedly occurring neighbourhood of a gene. Nucleic Acids Res. 28, 3442–3444 (2000).

60. Raudvere, U. et al. g:Profiler: a web server for functional enrichment analysis and conversions of gene lists (2019 update). Nucleic Acids Res. 47, W191–W198 (2019).

61. Xu, S. et al. Using clusterProfiler to characterize multiomics data. Nat. Protoc. 19, 3292–3320 (2024).

